# Associative memory in human cancer cells

**DOI:** 10.1101/2025.09.29.679179

**Authors:** Stéphane Dayot, Jean-Philippe Morretton, Nadia Elkhatib, Severine Lecourt, Camille Lobry, Guillaume Montagnac

## Abstract

In psychology and neuroscience, associative memory refers to the capacity to learn and remember a link between two unrelated items. Although associative memory is widely believed to be restricted to animals possessing a complex nervous system, several reports have suggested that single-cell organisms can be conditioned to develop an associative memory-like behavior. Here, we report that human cancer cell lines can be conditioned to associate an extracellular matrix component and Gefitinib, a drug that reduces cell migration velocity. Collagen-I was periodically paired with Gefitinib and we observed that conditioned cells progressive decreased migration velocity on collagen-I but not on other extracellular matrix components. We identified the adenosine receptor ADORA2A as a key actor regulating the acquisition of associative memory. We also observed that the magnitude of the conditioned response oscillated over time with the same periodicity as paired stimuli presentations during conditioning. We found that mitochondria morphology oscillated with the same periodicity, suggesting that memory and energy metabolisms are linked. We propose that human cancer cells can be conditioned to integrate a link between two stimuli from their environment in a process that may allow to anticipate future stress exposition.

## INTRODUCTION

Associative memory is best illustrated by the seminal Pavlov experiment during which a dog conditioned to associate the sound of a bell together with food progressively developed a conditioned response (*1*). Several bell/food paired presentations are required before the dog starts to salivate in response to the bell alone. The current mechanistic understanding supporting associative memory involves synaptic plasticity and the progressive reinforcement of connections between neurons that fire together (*2*). Accordingly, associative memory is widely considered to be the privilege of organisms that developed a nervous system. However, beyond neuronal networks, mathematical models and simulations suggest that proteins and/or nucleic acid-based networks such as gene regulatory networks can theoretically support associative memory (*3–5*). In addition, studies in unicellular organisms have reported behaviors that are highly reminiscent of associative memory (*6–9*). Although controversial, associative memory in single-cell organisms would certainly represent a competitive advantage in complex environments. It is less clear how a cellular associative memory would integrate into the developmental and homeostatic programs of multicellular organisms. To our knowledge, only one study investigated the possibility of associative memory in mammalian cells (*10*). The authors did not find any evidence of conditioning in human macrophages, but issues regarding the design of the conditioning protocol do not allow to exclude the possibility of associative memory in mammalian cells. Beyond physiology, cancer cells are widely recognized for their plasticity and their capacity to adapt to treatments (*11*). In this context, learning capabilities may represent an advantage to thrive in challenging environments. Here, we set out to determine if human cancer cell lines can perform associative memory.

## RESULTS

To test the possibility of associative memory in breast cancer MDA-MB-231 cells, we developed a protocol that is adapted to mammalian cell culture and that matches the requirements for associative conditioning. We chose to use Collagen-I as an environmental stimulus and Gefitinib as a stress stimulus. Collagen-I is an extracellular matrix protein that is not present in the complete medium in which cells are grown. Gefitinib is an inhibitor of the tyrosine kinase activity of the epidermal growth factor receptor (EGFR). The conditioned response we chose to follow is a reduction of cell migration velocity. This choice was based on observations that MDA-MB-231 cells plated on Col-I-coated glass migrated fast (Supp. Fig. 1A), and that Gefitinib reduced migration velocity (Supp. Fig. 1B). We also observed that Gefitinib induced an enlargement of focal adhesions marked with vinculin (Supp. Fig. 1C and D). Enlarged focal adhesions are probably linked to reduced cell velocity and these two parameters were hereafter considered as features of the response to Gefitinib.

Our conditioning protocol consisted in repeated cycles of cell culture in two sequential environments. In the first sequence, MDA-MB-231 cells were seeded on Collagen-I-coated glass in complete medium for 2 hours before 10 µM Gefitinib was added in the medium. Cells were kept on this environment for 3 days. In the second sequence, cells on the first environment were harvested and transferred onto naked glass in complete medium. Cells were kept for 3 days in this second environment where both Collagen-I and Gefitinib are absent (Figure 1A). Such a 6 days cycle was repeated up to 5 times. At the end of each cycle, control and conditioned cells were transferred onto Collagen-I-coated glass (without Gefitinib) and cell velocity was measured by tracking individual cells migrating over a 16h period. We observed that velocity was constant in control cells but that it progressively decreased with the accumulation of cycles in conditioned cells (Figure 1B). This new response to Collagen-I was the strongest after 4 cycles (no further decrease after 5 cycles). Thus, cells were conditioned for 4 cycles in subsequent experiments. The new response was specific of Collagen-I as conditioned cells plated on naked glass or on Collagen-IV-coated glass migrated as fast as their respective controls (Figure 1C). Accordingly, conditioned cells seeded on collagen-I showed enlarged focal adhesions as compared to control cells and to conditioned cells seeded on collagen-IV (Supp. Fig. 2A and B). We next conditioned cells using different control protocols (Figure 1D). Periodic presentation of one or the other stimulus alone for a period equivalent to four training cycles did not modulate cell velocity on collagen-I as compared to control cells (Figure 1D and E). Similarly, continuous presentation of one of the stimuli with alternating presentation of the other one, or continuous presentation of both stimuli together, did not generate a specific response (Figure 1D and E). Finally, alternating Gefitinib and collagen-I exposition periods did not modulate the response to collagen-I either (Figure 1D and E). Thus, strict pairing of the two stimuli during the conditioning protocol is required for cells to develop a new response to collagen-I. Collectively, these results demonstrate that MDA-MB-231 cells are endowed with associative memory capability. We also tested associative memory in lung cancer PC-9 cells. These cells bear a mutation in EGFR that makes them very sensitive to Gefitinib treatment and we thus chose to work at a concentration of 10 nM (*12*). We observed that PC-9 cells conditioned for 4 cycles showed a significant reduction in migration velocity on collagen-I but not on glass or on collagen-IV (Supp. Fig. 2C and D). These results suggest that PC-9 cells can also perform associative memory. We next aimed to test whether our results were specific of the collagen-I/Gefitinib association or whether other associations are possible. Using a variation of the protocol described above, we paired collagen-IV together with Gefitinib and observed that conditioned cells migrated significantly slower than control cells when seeded on collagen-IV but not on collagen-I (Figure 1F and G). These results show that MDA-MB-231 cells can integrate a specific link between different components of their environment.

**Figure 1.**
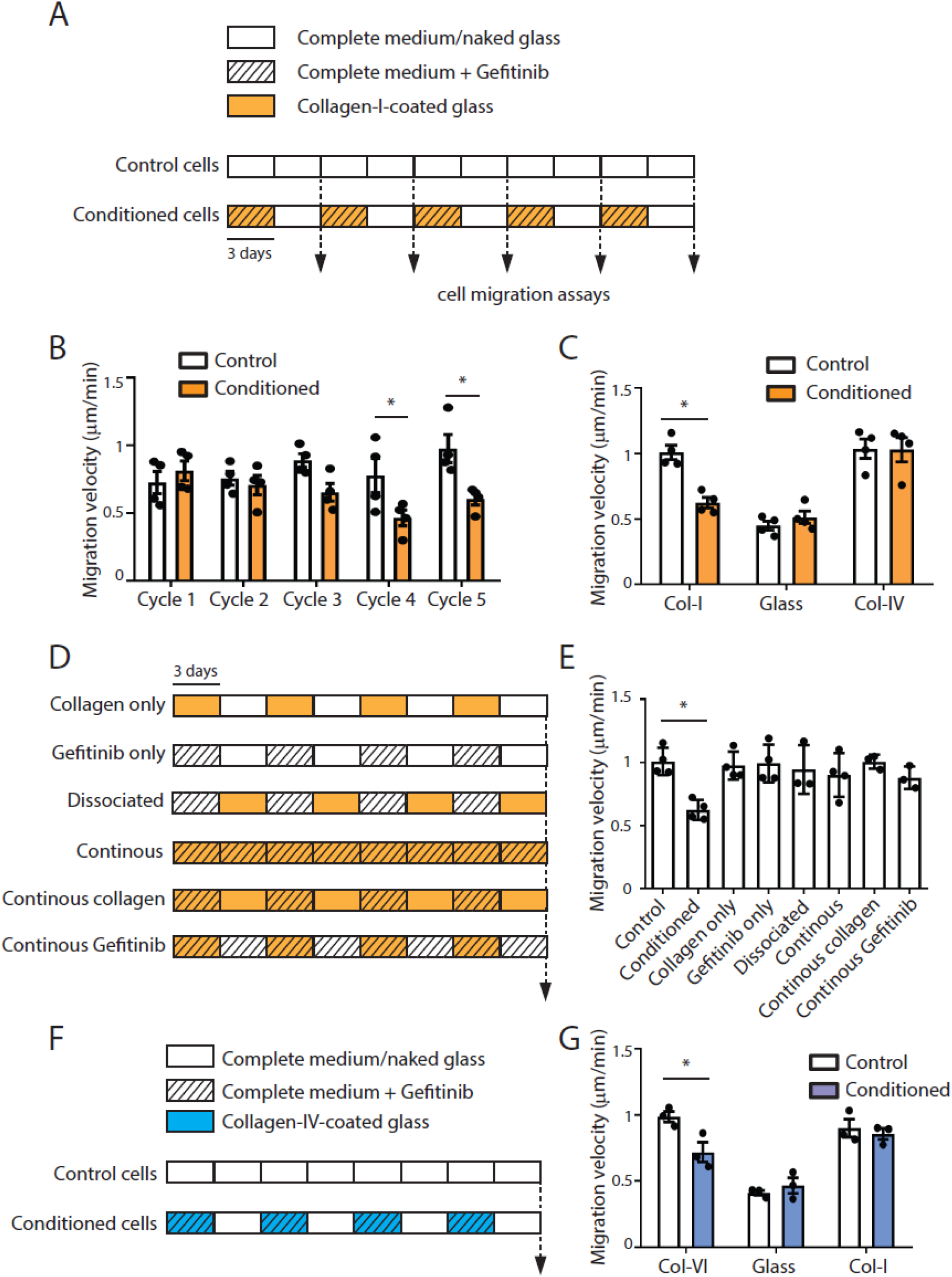
Evidence of conditioning in MDA-MB-231 cells. **a**, Description of the culture conditions for the conditioning protocol using the collagen-I/Gefitinib association and for control cells. Cells were split every 3 days (vertical bars). **b**, MDA-MB-231 cells conditioned for the indicated number of cycles as in a were seeded on collagen-I-coated glass and imaged every 10 minutes for 16h. Results are expressed as the mean migration velocity in um/min (*P<0.001, One Way Analysis of Variance – ANOVA. N=4). **c**, MDA-MB-231 cells conditioned for 4 cycles were seeded on collagen-I-or collagen-IV-coated glass or on naked glass, as indicated, and imaged every 10 minutes for 16h. Results are expressed as the mean migration velocity in um/min (*P<0.001, One Way Analysis of Variance – ANOVA. N=4). **d**, Description of the culture conditions for the different control protocols. **e**, MDA-MB-231 cells conditioned for 4 cycles as in d were seeded on collagen-I-coated glass and imaged every 10 minutes for 16h. Results are expressed as the mean migration velocity in um/min (*P<0.001, One Way Analysis of Variance – ANOVA. N=4). **f**, Description of the culture conditions for the conditioning protocol using the collagen-IV/Gefitinib association and for control cells. **g**, MDA-MB-231 cells conditioned for 4 cycles as in f were seeded on collagen-I- or collagen-IV- coated glass or on naked glass, as indicated, and imaged every 10 minutes for 16h. Results are expressed as the mean migration velocity in um/min (*P<0.001, One Way Analysis of Variance – ANOVA. N=3). All results are expressed as mean ± SD.

We next aimed to address the mechanisms of cellular associative memory. We first reasoned that our conditioning protocol may result in a clonal selection. To address whether cellular conditioning is mediated through a global population response to the training or to oligoclonal selection of pre-existing adapted subclones we performed a cellular barcoding experiment. Cells were transduced with the LARRY barcoding library to contain a single barcode in their genomic DNA. Transduced cells were then subsampled (parental cell population) and submitted to the conditioning protocol or to the control protocol as in Figure 1A. Genomic DNA from the two populations was subsequently extracted and sequenced to identify barcodes present in the cellular populations. A strong selective pressure inducing oligoclonal selection should result in a stiff decrease of barcode diversity whereas a global population response mediated by all the clones should result in a globally conserved barcode diversity. To measure this diversity and richness of barcodes we calculated Shannon and Simpson indexes. Both indexes showed no significant variations between conditioned and control cell populations (Figure 2A and B). These results show that the conditioning protocol doesn’t induce an oligoclonal selection and maintain overall cellular diversity in the population.

**Figure 2.**
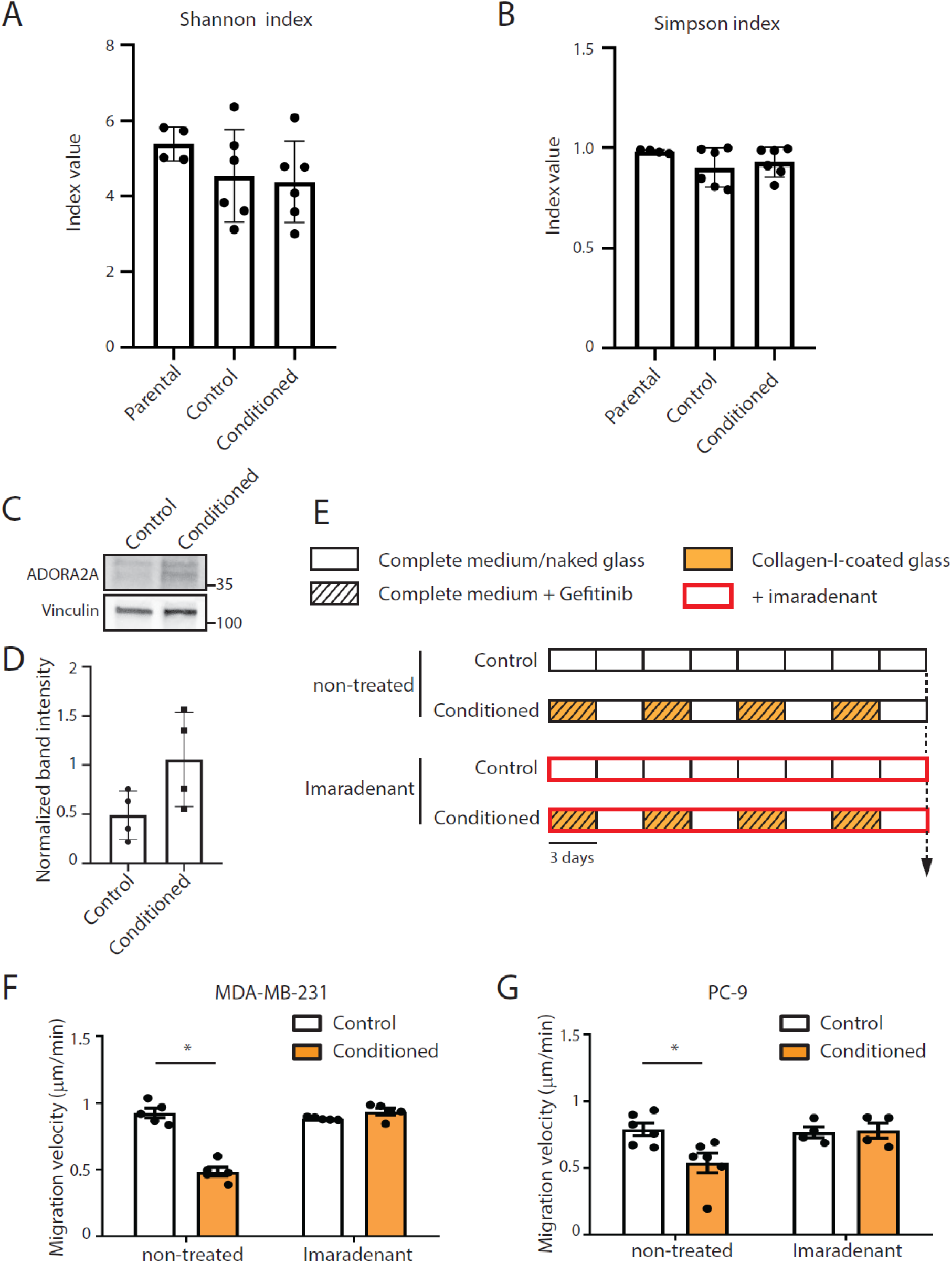
ADORA2A regulates cellular associative memory. **a**, **b,** Shannon index (a) and Simpson index (b) calculated from the barcoding experiment for Parental MDA-MB-231 cells and cells exposed to the conditioning or control protocol as indicated. Data are from two independent experiments performed in triplicates. **c**, Western blot analysis of ADORA2A expression in control and conditioned MDA-MB-231 cells. Vinculin was used as a loading control. **d**, quantification of ADORA2A band intensity as in c. **e**, Description of the culture conditions for the conditioning protocol using the collagen-I/Gefitinib association and for control cells, in the presence of imaradenant. Cells were split every 3 days (vertical bars). **f**, MDA-MB-231 cells conditioned for four cycles as in d were seeded on collagen-I-coated glass and imaged every 10 minutes for 16h. Results are expressed as the mean migration velocity in um/min (*P<0.001, One Way Analysis of Variance – ANOVA. N=5). **g**, PC-9 cells conditioned for four cycles as in d were seeded on collagen-I-coated glass and imaged every 10 minutes for 16h. Results are expressed as the mean migration velocity in um/min (*P<0.001, One Way Analysis of Variance – ANOVA. N=4).

We next compared the transcriptome of control MDA-MB-231 cells versus cells conditioned with the collagen-I/Gefitinib association. We observed an enrichment in pathways related to inflammatory response and to G protein-coupled receptors signaling in conditioned cells (Supp. Fig. 3A). In particular, the adenosine A2a receptor (ADORA2A) was found to be overexpressed in conditioned cells (Supp. Fig. 3B) and has been proposed to regulate the migration of some cancer cells (*13*). We first validated that ADORA2A protein is overexpressed in conditioned cells (Figure 2C and D). We observed that inhibition of ADORA2A with the specific inhibitor imaradenant did not modulate cell migration velocity nor focal adhesions size in control MDA-MB-231 cells seeded on collagen-I (Supp. Fig. 3C-E). In addition, imaradenant did not modulate the consequences of Gefitinib treatment on migration velocity and on focal adhesion enlargement in control cells (Supp. Fig. 3C-E). However, we observed that inhibiting ADORA2A during conditioning abolished the acquisition of the associative response as cells conditioned in the presence of the drug migrated as fast as control cells when tested on collagen-I (Figure 2E and F). We obtained similar results when using PC-9 cells (Figure 2E and G). Thus, ADORA2A activity is required for the formation of associative memory in human cancer cells.

We next aimed to test for how long the memory was maintained after the end of conditioning. For this, MDA-MB-231 cells conditioned for 4 cycles with a collagen-I/Gefitinib association were tested on collagen-I at different time points just before and after the end of conditioning. In the meantime, cells were kept on naked glass, in the absence of Gefitinib (Figure 3A). We observed that cell velocity was low at the end of the conditioning protocol (day 0) but progressively increased over the next 3 days to reach the same velocity as control cells (Figure 3B). This suggested that the conditioned response vanishes very quickly after the end of conditioning. However, we observed that cell velocity decreased during the following 3 days and then increased again during the next 3 days (Figure 3B). Migration speed variations over time were not observed in conditioned cells tested on collagen-IV (Figure 3B). Thus, the associative response oscillates over time with a periodicity of 6 days that matches the periodicity of stimuli co-presentation during the conditioning period. This oscillation was still detectable 1 month after the end of the conditioning protocol (Figure 3B). The periodicity was highly conserved, with time points of maximal conditioned response corresponding to the precise timing of Gefitinib presentation during the conditioning protocol (Figure 3B). We also observed that focal adhesion area increased on collagen-I, but not on collagen-IV, on day 0 post-conditioning (the time of maximal conditioned response) compared to 3 days after the end of conditioning, when no conditioned response was observed (Figure 3C and D). These data show that conditioned cells autonomously change their phenotype at regular intervals, specifically in response to collagen-I. Our results demonstrate that cellular associative memory oscillates, with the conditioned response being expressed only periodically.

**Figure 3.**
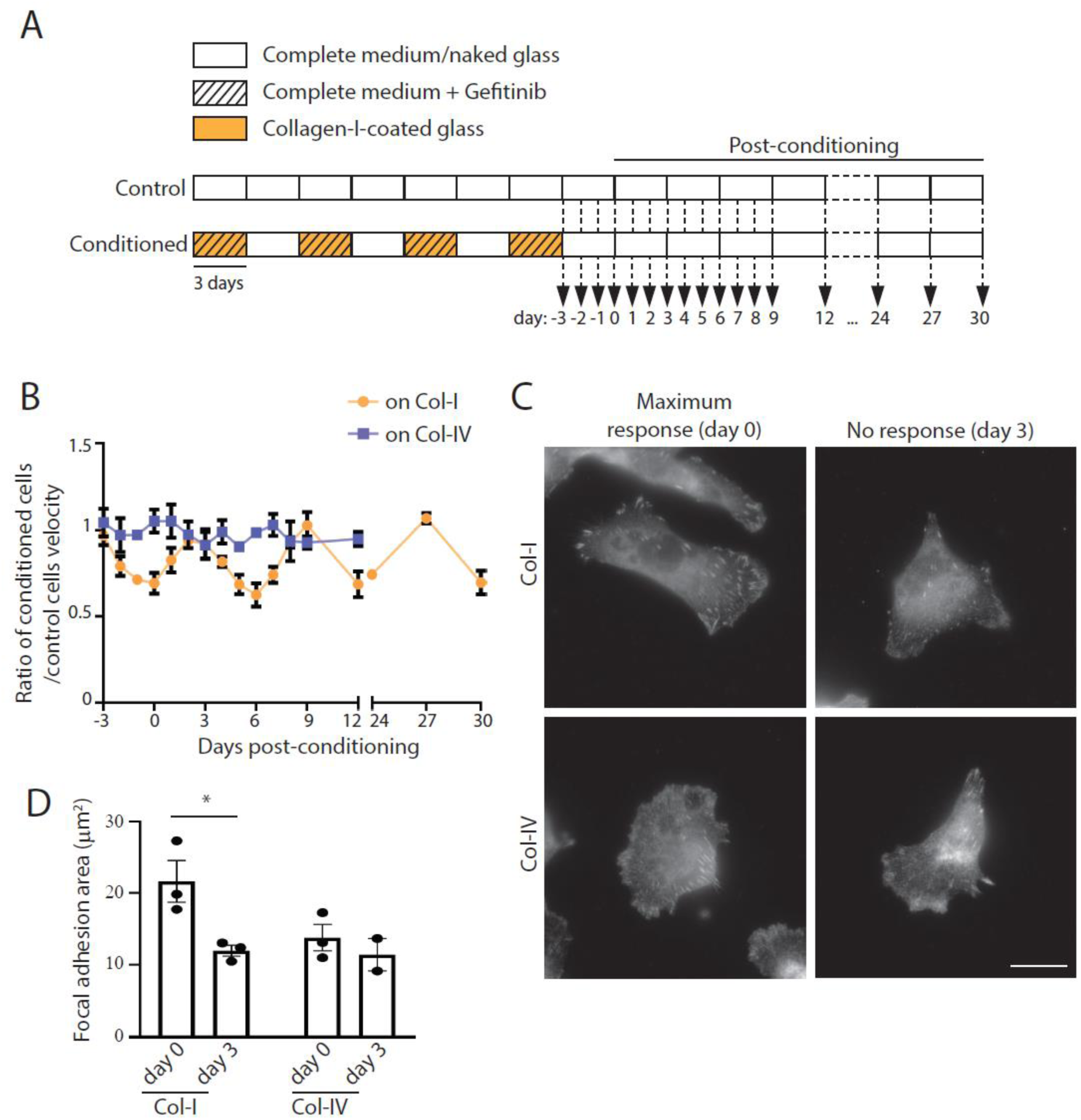
Cellular associative memory oscillates in time. **a**, Description of the culture conditions for the conditioning and post-conditioning protocol using the collagen-I/Gefitinib association and for control cells. Cells were split every 3 days (vertical bars). Arrows indicate time points (days post-conditioning) at which cells were transferred onto collagen-coated glass and imaged to measure cell migration velocity **b**, MDA-MB-231 cells conditioned for four cycles and then kept on naked glass for the indicated number of days as in a were seeded on collagen-I- or collagen-IV-coated glass and imaged every 10 minutes for 16h. Results are expressed as the mean migration velocity in um/min. **c**, Conditioned MDA-MB-231 cells seeded on collagen-I-coated glass (upper panels) or on collagen-IV-coated glass (lower panels) at day 0 or day 3 post-conditioning, as indicated, were fixed and stained for vinculin. Scale bar: 10 μm. **d**, Quantification of the total area occupied by vinculin positive focal adhesions as in c (* P<0.001, One Way Analysis of Variance – ANOVA. N=3). Results are expressed as mean ± SD.

We observed that ADORA2A activity is required during conditioning for cells to integrate the collagen-I/Gefitinib association. We next asked whether ADORA2A was also required to maintain the memory after the end of conditioning. We observed that cells treated with imaradenant for 6 days post-conditioning did not exhibit the associative response, in contrast to control conditioned cells, which migrated more slowly on collagen-I on day 6 post-conditioning (Supp. Fig. 4A and B). In addition, ADORA2A depletion using specific siRNAs delivered at day 0 and day 3 post-conditioning also resulted in the abrogation of the conditioned response normally observed at day 6 (Supp. Fig. 4C and D). Thus, ADORA2A is required to generate and maintain the periodic phenotypic oscillation that characterizes cellular associative memory.

To better characterize the changes associated with this phenotypic oscillation, we performed RNA sequencing (RNA-seq) analyzes comparing conditioned cells at time points of maximal (day 0 post-conditioning) and no (day 3 post-conditioning) conditioned response. Gene set enrichment analyzes showed an enrichment in pathways associated with mitochondrial functions (Figure 4A). Gefitinib has been shown to instigate mitochondrial dysfunctions through the production of reactive oxygen species (*14*). Indeed, we observed that Gefitinib treatment induced an increased mitochondrial area together with an increased filamentous morphology characterized by longer mitochondria branches (Supp. Fig. 5A-C). Upon conditioning, we observed that cells similarly displayed an increased mitochondria network area together with longer branches as compared to control cells when tested on collagen-I (Figure 4B-D). This mitochondria fusion phenotype was observed at day 0 post-conditioning (time point of maximal conditioned response) but not at day 3 post-conditioning (time point of no response) suggesting that mitochondria morphology oscillates as well in conditioned cells (Figure 4B-D). Thus, mitochondria dynamics oscillates together with migration capabilities in conditioned cells. Because ADORA2A controls the acquisition of the conditioned response, we next asked whether it also controlled mitochondria dynamics. Inhibiting ADORA2A with imaradenant or with specific siRNAs for 3 days before the end of the conditioning period was sufficient to rescue mitochondria morphology in conditioned cells (Figure 4E and F). Thus, ADORA2A regulates both migration velocity and mitochondria oscillation in conditioned cells.

**Figure 4.**
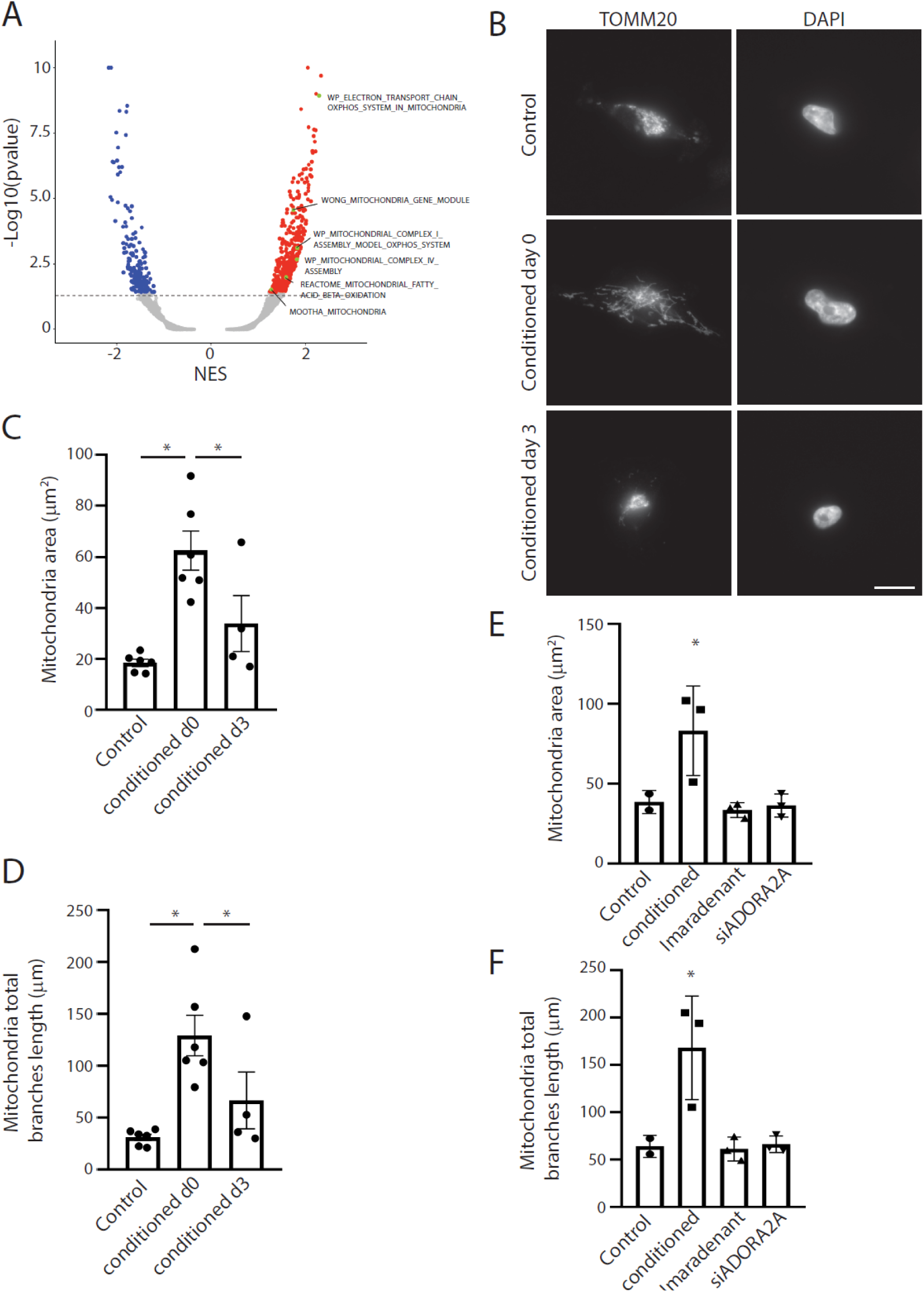
Mitochondria dynamics oscillates in conditioned cells. **a**, Volcano plot representing gene set enrichment analysis (GSEA) results of differentially expressed genes between conditioned MDA-MB-231 cells at day 0 versus at day 3 post-conditioning. **b**, Control or conditioned MDA-MB-231 cells at time points of maximal (day 0) or no-response (day 3) and seeded on collagen-I were fixed and stained for Tomm20. Scale bar: 10 μm. **c-d**, Quantification of total mitochondria area (c) and of the total length of branches (d) in cells as in b (* P<0.001, One Way Analysis of Variance – ANOVA. N=4). **e-f**, Conditioned MDA-MB-231 cells were treated or not for 3 days with imaradenant or with ADORA2A specific siRNAs, as indicated, just before the end of conditioning. Cells were then seeded on collagen-I before to be fixed and stained for Tomm20 in order to measure total mitochondria area (e) and the total length of branches (f) (* P<0.001, One Way Analysis of Variance – ANOVA. N=3).

## DISCUSSION

Overall, we report that cancer cells show associative memory capability as they can integrate a link between an extracellular matrix components (collagen-I or collagen-IV) and a drug. Within the frame of the protocol used in this study, we characterized the oscillatory nature of cellular associative memory and identified the adenosine receptor ADORA2A as a key actor in this process. ADORA2A has been shown to alleviate mitochondria injury induced by reactive oxygen species (*15*), dysfunctions that are promoted by Gefitinib (*14*). A potential feedback loop between Gefitinib and ADORA2A actions on mitochondria may set the oscillatory nature of the conditioned response. The entanglement of mitochondria dynamics with the oscillatory pattern of cellular associative memory also suggests that energy metabolism may underline parts of the learning mechanism. However, the exact mechanism triggering the conditioned response remains to be investigated. Overall, we report that human cancer cells are endowed with associative memory capabilities that may allow them to anticipate stressful conditions periodically arising in their environment.

## METHODS

### Cell lines and extracellular matrices

MDA-MB-231 cells (a gift from P. Chavrier, Institut Curie, Paris, France; ATCC HTB-26) and PC-9 cells (a gift from F. André (Inserm U981), Gustave Roussy, Villejuif, France; RRID:CVCL_B260) were grown in DMEM Glutamax supplemented with 10% foetal calf serum at 37°C in 5% CO2. All cell lines have been tested for mycoplasma contaminations. For experimental protocol and cell migration assay, cells were plated on glass coated with a 50 µg/ml solution of collagen-I (Thermo Fisher Scientific - Cat. Nr. A10483-01) or a 20 µg/mL solution of Collagen IV (Corning – Cat. Nr. 354233).

### Conditioning protocol

MDA-MB-231 and PC-9 cells were grown on Collagen-I-coated glass in complete medium for 1 hours before 10 µM (MDA-MB 231) or 10 nM (PC-9) Gefitinib was added in the medium. Cells were kept on this environment for 3 days. Cells were then harvested and transferred onto naked glass in complete medium without Collagen-I or Gefitinib. Cells were kept for 3 days in this second environment and this 6 days cycle was repeated up to 5 times. At the end of each cycles, cells were harvested and tested for cell migration assay.

### *Time-lapse micros*c*opy and cell migration velocity quantification*

Time-lapse videos were acquired on a Leica epifluorescence microscope (Leica Microsystems Ltd., Wetzlar, Germany) through 10x (Fluotar N.A. 0.32 PH1) objective. A PeCon chamber i8 was installed on the microscope to perform live imaging at 37°C with 5% CO2. The microscope was steered by the Leica dedicated LasX software®. Control and conditioned cells were harvested and seeded on a 96 glass-bottom plate coated or not with collagen I or collagen IV.

Cells were then allowed to adhere for 1-2 h and imaged in bright field or phase contrast, at a 1 image every 20min for 16h. Cells were manually tracked using the “Manual tracking” plugin of FIJI. Migration velocity were obtained with the Ibidi “Chemotaxis and Migration Tool” in FIJI. For each condition, 50-80 cells were manually tracked from at least 3 independent experiments. Data are expressed as mean ± SEM.

### Antibodies and drugs

Mouse monoclonal anti-Vinculin (clone VIIF9 – Cat. Nr. MAB3574) antibody and mouse polyclonal anti-ADORA2A (A2AR – Cat. Nr. SAB1408884) were obtained from Sigma-Aldrich and mouse monoclonal anti-Tomm20 (F-10 – Cat. Nr. sc-17764) antibody was obtained from Santa Cruz Biotechnology Inc. (Santa Cruz, CA, USA). Antibodies used for western blot analyses were diluted at 1:1000 in PBS-0.1% Tween-5% BSA or 5% non-fat dried milk. For immunofluorescences, antibodies were diluted 1:200 in PBS-0.3% BSA. HRP-conjugated anti-mouse and anti-rabbit antibodies for western blot were from Jackson ImmunoResearch Laboratories (West Grove, PA, USA) and were used at a dilution of 1:3000. Alexa-conjugated antibodies were from Molecular Probes (Invitrogen) and were used at a dilution of 1:200. Rat tail Collagen-I (Cat. Nr. A10483-01) was purchased from GIBCO and Mouse Collagen-IV was purchased from Corning. Gefitinib (Cat. Nr. HY-50895) and Imaradenant (AZD4635 – Cat. Nr. HY-101980) were purchased from CliniSciences. Gefitinib was used at a final concentration of 10 μM for MDA-MB-231 conditioning and 10 nM for PC-9 cells. Imaradenant was used at a final concentration of 2 nM.

### Western Blots

For Western Blot experiments, cells were lysed in ice cold RIPA buffer (500mM Tris-HCl, pH 7.4, 150 mM NaCl, 0.25% deoxycholic acid, 1% NP-40, 1mM EDTA.) supplemented with protease and phosphatase inhibitors. Protein concentration was measured using Pierce™ Coomassie Plus (Bradford) Assay Kit (1856210 according to the manufacturer’s instructions in order to load equal amount of proteins. Antibodies were diluted at 1:1000 in PBS - 0.1% Tween - 5% BSA or 5% non-fat dried milk. Analyzes of bands densitometry were performed using ImageJ.

### RNA interference

For siRNA depletion, cells were plated on naked glass wells of a 12 well plates at 30% confluence. After 24 h, cells were treated with the indicated siRNA (30 nM) using RNAimax (Invitrogen, Carlsbad, CA) according to the manufacturer’s instruction. Protein depletion was maximal after 72 h of siRNA treatment as shown by immunoblotting analysis with specific antibodies. The following siRNAs were used: ADORA2A (1), 5’- CAUGCUGGGUGUCUAUUUG-3’; ADORA2A (2), 5’-GGAGUGUCCUGAUGAUUCA-3’; non-targeting siRNAs (siControl), ON-TARGETplus Non-Targeting SMARTpool siRNAs (Dharmacon D-001810-01).

### Immunofluorescence microscopy and fluorescence quantification

MDA-MB-231 were seeded on naked or Collagen I or IV-coated 12mm coverslips and incubated at 37°C with 5% CO2 for 12h. Cells were fixed in 4% paraformaldehyde (PFA - ThermoFischer) for 15min at RT before being washed with PBS. Cells were incubated with primary antibodies (1:200) for 1h at RT before being washed with PBS. Finally, cells were incubated with secondary antibodies (1:200) for 1h at RT and processed for immunofluorescence microscopy. Cells were imaged through a 63× 1.40NA UPL APO objective lens of a DMi8 Thunder imager microscope (Leica) equipped with an DFC9000 sCMOS camera (Leica) and steered by LAS X software (Leica).

To analyse focal adhesions (FAs), Vinculin-positive structures were segmented in ImageJ following step-by-step protocol from Horzum U paper (MethodX) excluding objects with area smaller than 250 nm. Particles were analysed and the total area were measured / cells. Total area of FAs were measured from a number of cells ranging from 10 to 60 for each condition and from 3 independent experiments. Data are expressed as mean ± SEM. Mitochondria morphology was analysed using Mitochondria Analyser from FiJi Plugin. Tom-20-positive structures were segmented using 2D Threshold tool from the plugin and with the following parameters : “Substract Background : Rolling 1.25 microns” ; “Radius : 0.97” ; “Max Slope : 1.80” “Gamma : 0.80” ; “Threshold Method : Median” ; “Block Size : 1.250” ; “C-Value : 5” with “Despeckle” and Outliers removed with Radius below 1 pixel. Mitochondrial area and Total branches length were extracted from the 2D Analysis tool from the plugin and were measured from at least 30 cells for each conditions and from 3 independent experiments. Data are expressed as mean ± SEM.

### Barcoding experiment

MDA-MB-231 cells were infected with the LARRY (Lineage and RNA Recovery) barcode lentiviral library (*16*) (Addgene plasmid #140024, Camargo Lab). This library clonally tags cells with unique DNA barcodes to enable lineage tracing.

After infection and cell sorting, cells were grown following the same conditioning protocol than before and then harvested at the end of cycle 5 for DNA extraction. Cells were lysed to extract genomic DNA using AllPrep DNA/RNA Micro Kit (Qiagen 80284). Barcode regions were amplified using primers following the LARRY genomic protocol to specifically target barcode sequences integrated in the genome (*16*). Amplified barcode libraries were sequenced on Illumina NextSeq 500 sequencer

### Barcoding experiment analysis

Raw reads were aligned to the reference LARRY barcode library using Bowtie2 (*17*) with -- very-fast option. Aligned reads were processed with Mageck count (*18*) to generate raw count matrices of barcode abundances per sample.

Barcode whose coverage where >50 reads in more than 2 samples per group where kept for downstream analysis.

Diversity of barcodes among samples and group was measured using Shannon and Simpson indices. Shannon diversity indices (H^′^) were calculated per sample from filtered raw barcode counts according to:

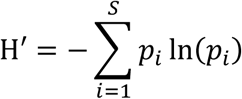

where 𝑝𝑖 denotes the relative abundance of barcode i and S the total number of barcodes in the dataset. Confidence intervals were calculated using classical variance formulas.

Simpson indexes were calculated per sample from filtered raw barcode counts according to :

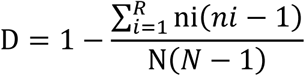

Where ni is the number of individuals of species i, N is the total number of individuals of all species and R is the richness, the total number of barcodes in the dataset.

Summary statistics were visualized using box plots with confidence intervals, and individual sample points were overlaid to represent variability. All computational analyses were performed in R (v4.3.3) using packages vegan, ecolTest, matrixStats, ggplot2, and dplyr.

### RNA sequencing and analysis

MDA-MB 231 cells were grown during 5 cycles of conditioning protocol and control and conditioned (days 0 post-conditioning) cells were harvested and seeded on collagen I-coated glass in a 12 well plate during 12 hours. For conditioned cells with no conditioning response (day 3 post-conditioning), cells were kept 3 more days on naked glass before to be seeded on collagen I-coated glass for 12 hours. RNA-seq was performed on poly(A) selected RNA extracted from the whole population of MDA-MB 231 seeded on Collagen-I coated glass using the RNeasy Plus Mini kit (Qiagen) following manufacturer’s protocol. Poly-(A)-selected, first stranded Illumina libraries were prepared with a modified TruSeq protocol using dUTP method. Three biological replicates per conditions were prepared. Libraries were size selected using AMPure XP SPRI beads, were amplified by PCR (maximum 16 cycles) and purified with AMPure XP beads and paired-end sequenced (75 bp) on the Illumina NextSeq 500 sequencer. Sequencing reads were pseudo aligned to the ENSEMBL GRCm38 version of the human genome and abundance quantified using Kallisto. Differential expression analysis was done using Sleuth R package. Volcano plots and other were plotted using R packages ggplot2.

Gene set enrichment was performed using R package fgsea (v1.20.0) and MsigDB (v7.5.1) HALLMARK, C5 and C2 collections. Gene sets with less than 15 genes or with more than 500 genes were excluded from the analysis. GSEA was performed using weighted enrichment statistic on a pre-ranked list of genes. Genes were ranked using their log2 fold change of expression between compared conditions. Gene sets with an FDR ≤ 0.25 and a nominal Pvalue ≤ 0.05 were considered as significantly enriched. GSEA results were plotted using R package ggplot2.

### Statistical analyses

Statistical analyses have been performed using One Way Analysis of Variance (ANOVA) followed by All Pairwise Multiple Comparison Procedure (Tukey’s method) for figures 1E ; 2F ; 2G ; 4I ; 4J; Supp Fig 1A ; Supp Fig 1B ; Supp Fig 4B ; Supp Fig 4D. Two Way Analysis of Variance (ANOVA) followed by All Pairwise Multiple Comparison Procedure (Tukey’s method) have been performed for figure 1B ; 1C ; 1G ; 3E ; 4C ; 4D and Supp Fig 2B. Unpaired t-test comparison has been performed on Fig 3E ; Supp Fig 1D ; Supp Fig 5B and Supp Fig 5C. All data are presented as mean of at least three independent experiments ± SEM. All statistical analyses were performed using GraphPad Prism software.

## Data availability

RNAseq data are available at Gene Expression Omnibus database under accession code GSE308458.

## Acknowledgment

The authors wish to thank the imaging facility of Gustave Roussy (PFIC) for help with image acquisition. Core funding for this work was provided by the Gustave Roussy Institute and the Inserm and additional support was provided by grants from the Fondation pour la recherche medical (DEQ20180339205), from the Institut National pour la recherche contre le Cancer (PL- BIO 2018-136 and PL-BIO 2022-087) and from the Agence Nationale de la Recherche (ANR- 23-CE16-0007-02) to GM.

S.D designed and performed experiments, analysed results and wrote the manuscript. JP.M, N.E and S.L, performed experiments. C. L. designed experiments and performed RNA-seq bioinformatic analyses. G.M supervised the study, designed experiments and wrote the manuscript.

The authors declare no competing interests. Correspondence and requests for materials should be addressed to or to

---

**Supp. Figure 1.**
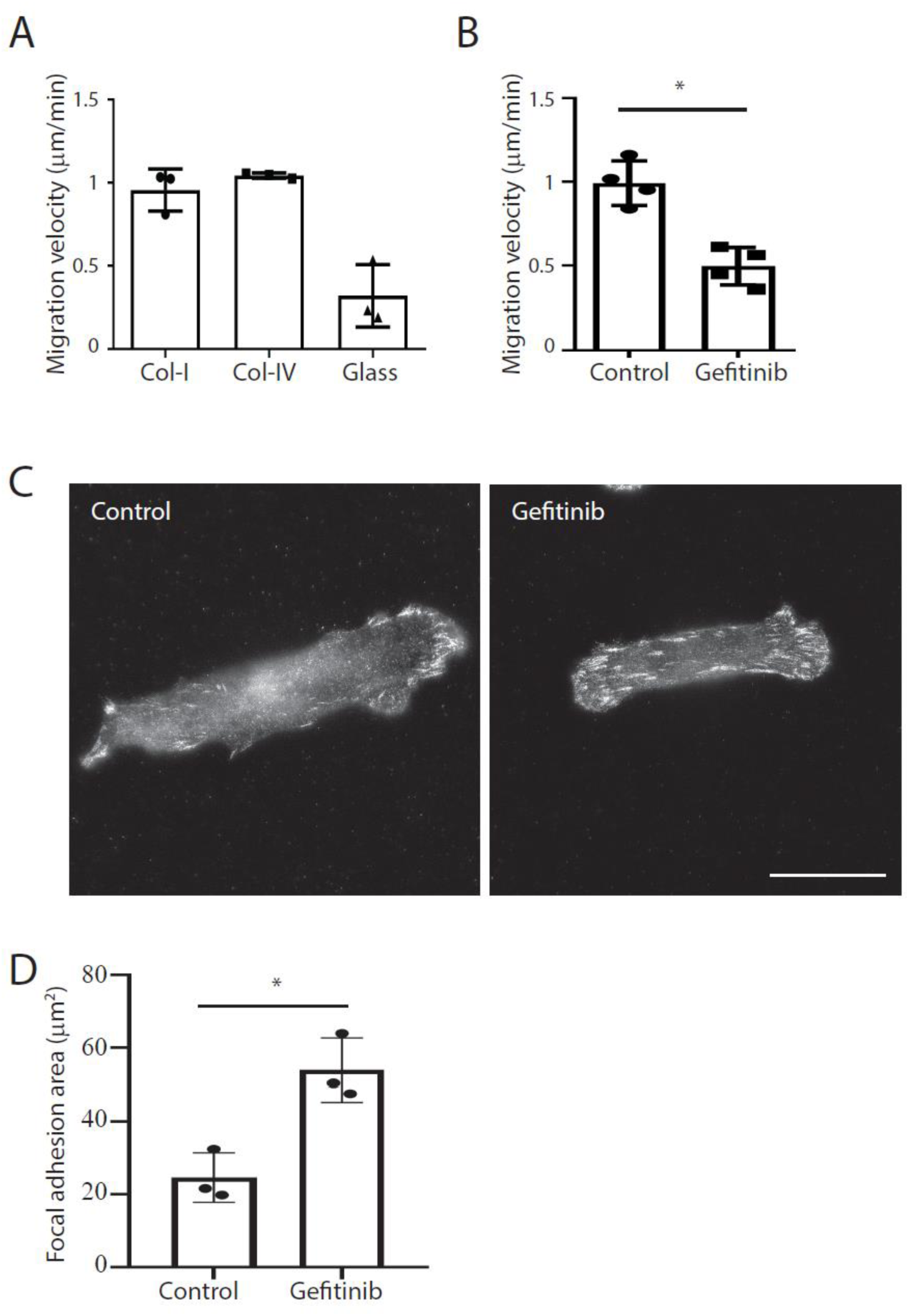
Characterization of cell response to collagen-I and to Gefitinib. **a**, MDA-MB-231 cells were seeded on naked glass or collagen-I- or collagen-IV-coated glass and imaged every 10 min for 16h. Results are expressed as the mean migration velocity in µm/min. **b**, MDA-MB-231 cells treated or not with Gefitinib and seeded on collagen-I were imaged every 10 min for 16h. Results are expressed as the mean migration velocity in µm/min (*P<0.001, Student’s t test. N=4). **c**, MDA-MB-231 cells seeded on collagen-I and treated or not with Gefitinib, as indicated, were fixed and stained for vinculin. Scale bar: 10 μm. **d**, Quantification of total focal adhesion area as in c (* P<0.001, Student’s t test. N=3).

**Supp. Figure 2.**
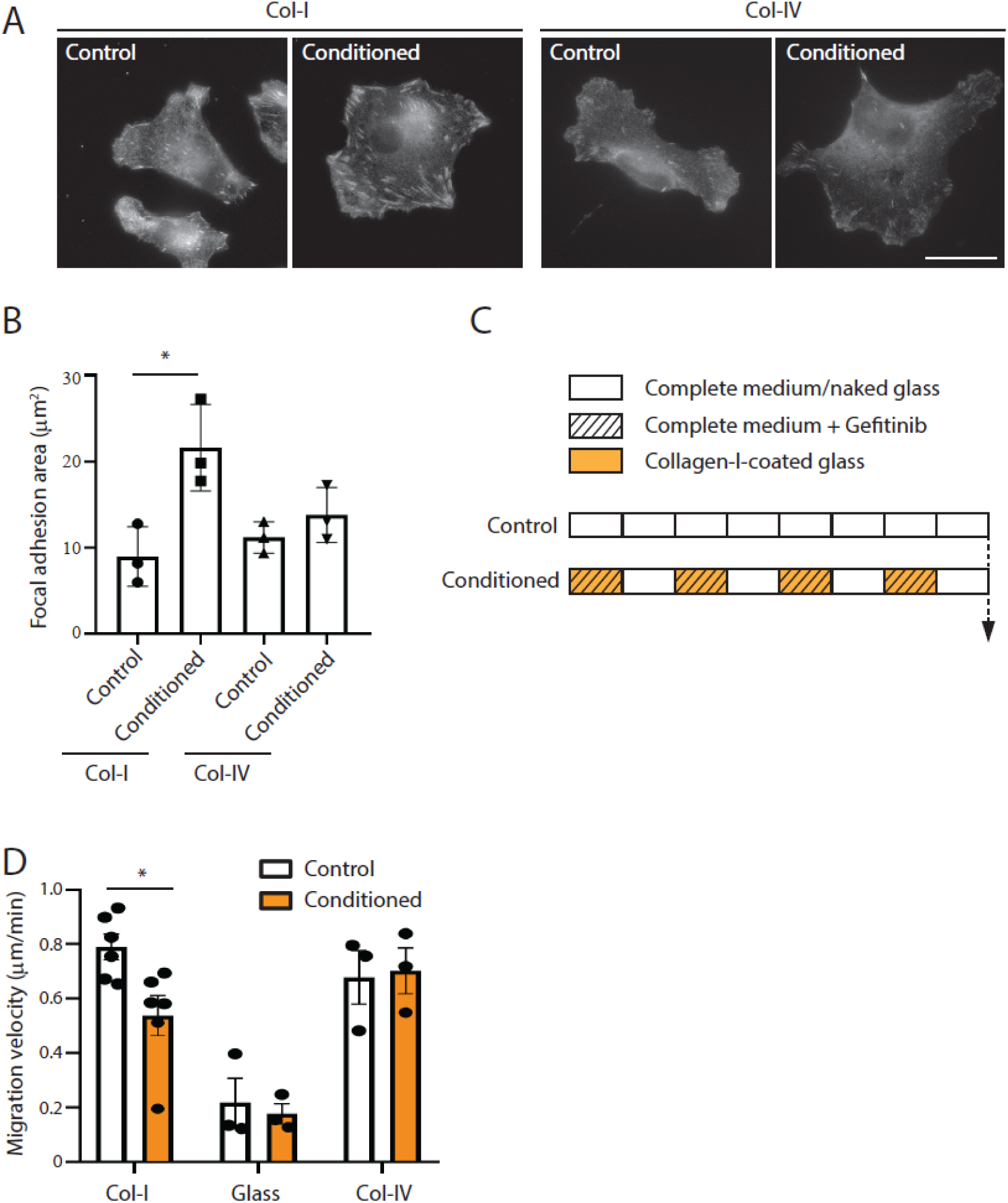
Characterization of the conditioned response in MDA-MB-231 and PC-9 cells. **a**, Control or conditioned MDA-MB-231 cells were seeded on collagen-I or collagen-IV, as indicated, before to be fixed and stained for vinculin. Scale bar: 10 μm. **d**, Quantification of total focal adhesion area as in a (* P<0.001, One Way Analysis of Variance – ANOVA. N=3). **c**, Description of PC-9 cells culture conditions for the conditioning protocol using the collagen-I/Gefitinib association and for control cells. Cells were split every 3 days (vertical bars). **d**, Control or conditioned PC-9 cells seeded on the indicated extracellular matrix, were imaged every 10 min for 16h. Results are expressed as the mean migration velocity in µm/min (*P<0.001, One Way Analysis of Variance – ANOVA. N=3).

**Supp. Figure 3.**
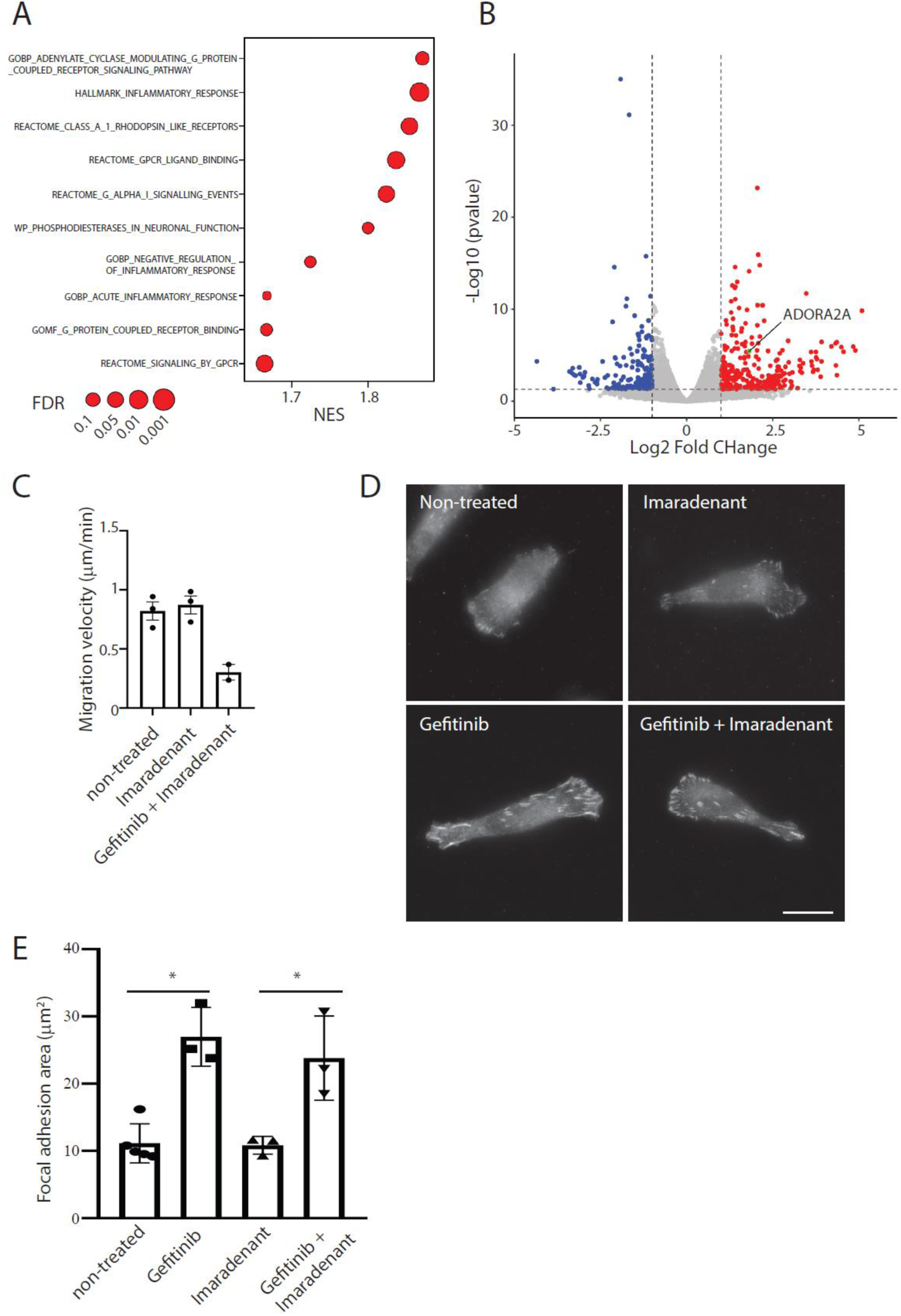
RNAseq and characterization of cell response to imaradenant. **a**, List of enriched pathways using Gene Set Enrichment Analysis comparing control and conditioned MDA-MB-231 cells. **b**, Volcano plot of differentially expressed genes between control and conditioned MDA-MB-231 cells. **c**, MDA-MB-231 cells treated or not with imaradenant and Gefitinib, as indicated, and seeded on collagen-I were imaged every 10 min for 16h. Results are expressed as the mean migration velocity in µm/min. **d**, MDA-MB-231 cells seeded on collagen-I and treated or not with Gefitinib and imaradenant, as indicated, were fixed and stained for vinculin. Scale bar: 10 μm. **e**, Quantification of total focal adhesion area as in b (* P<0.001, One Way Analysis of Variance – ANOVA. N=3).

**Supp. Figure 4.**
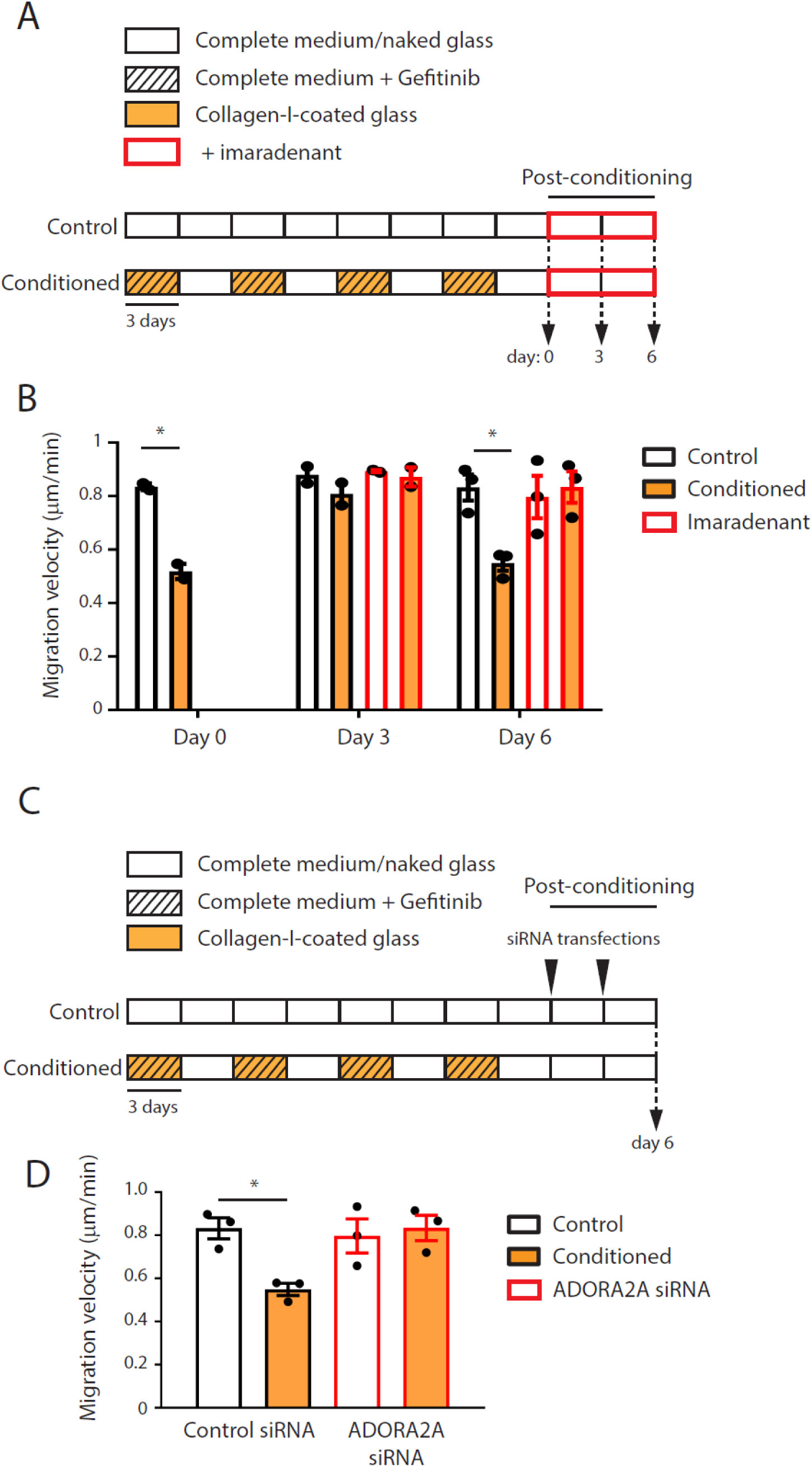
Role of ADORA2A in maintaining the conditioned response oscillation. **a**, Description of MDA-MB-231 cells culture conditions for the conditioning and post-conditioning protocol using the collagen-I/Gefitinib association and for control cells. Cells were split every 3 days (vertical bars). **b**, Control or conditioned MDA-MB-231 cells kept on naked glass for the indicated time post-conditioning and treated or not with imaradenant, as indicated, were seeded on collagen-I and imaged every 10 min for 16h. Results are expressed as the mean migration velocity in µm/min (* P<0.001, One Way Analysis of Variance – ANOVA. N=2). **c**, Description of MDA-MB-231 cells culture conditions for the conditioning protocol and post-conditioning treatment with ADORA2A specific siRNAs. Cells were split every 3 days (vertical bars). **d**, Control or conditioned MDA-MB-231 cells treated as in c were seeded on collagen-I and imaged every 10 min for 16h. Results are expressed as the mean migration velocity in µm/min (* P<0.001, One Way Analysis of Variance – ANOVA. N=3).

**Supp. Figure 5.**
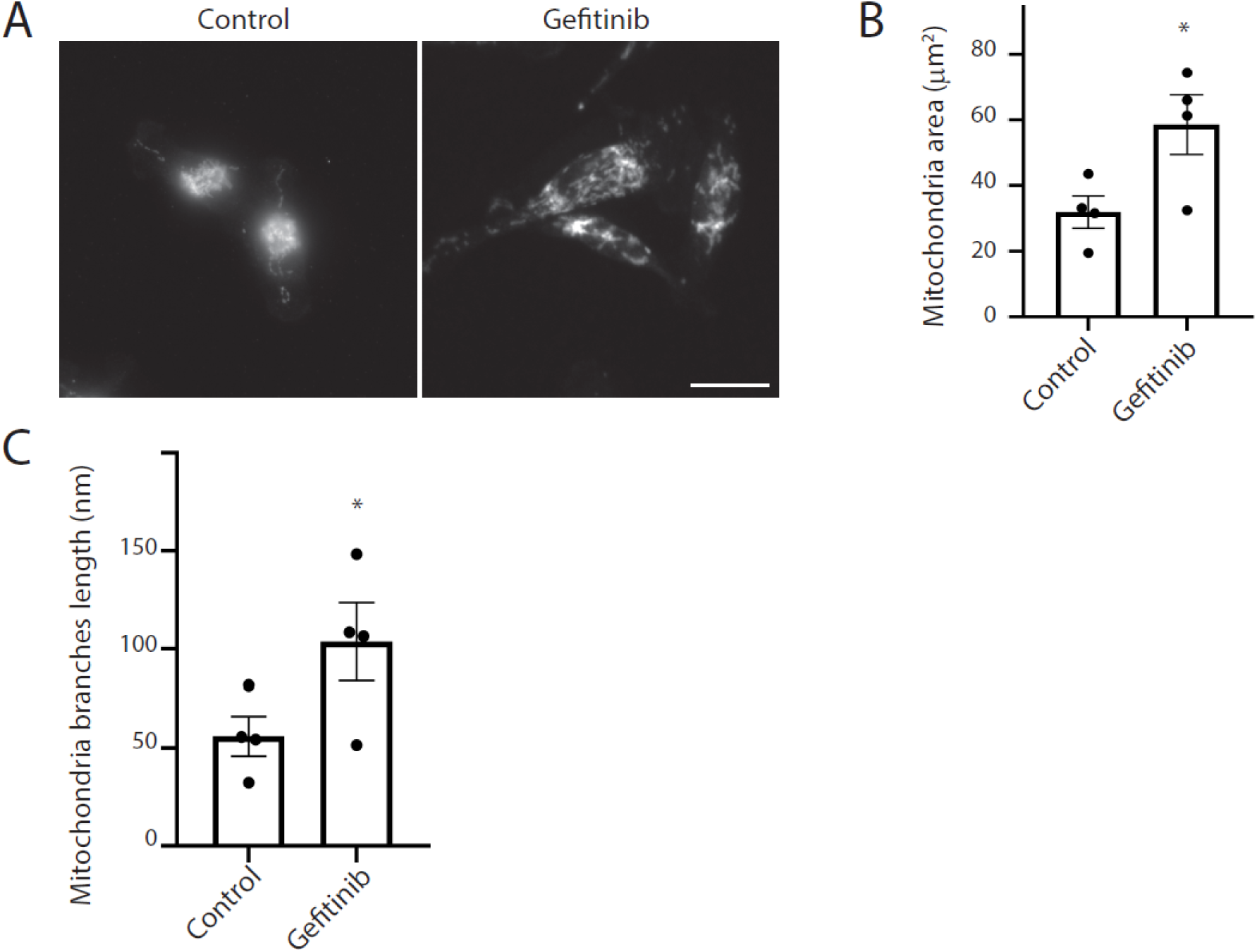
Characterization of mitochondria dynamics. **a**, Control MDA-MB-231 cells seeded on collagen-I and treated or not for 24h with Gefitinib were fixed and stained for Tomm20. Scale bar: 10 μm. b-c, Quantification of total mitochondria area (b) and of the total length of branches (c) in cells as in a (* P<0.001, Student’s t test. N=4).

