## Supplemental Material for "Associative memory in human cancer cells"

### Supplementary Figures:

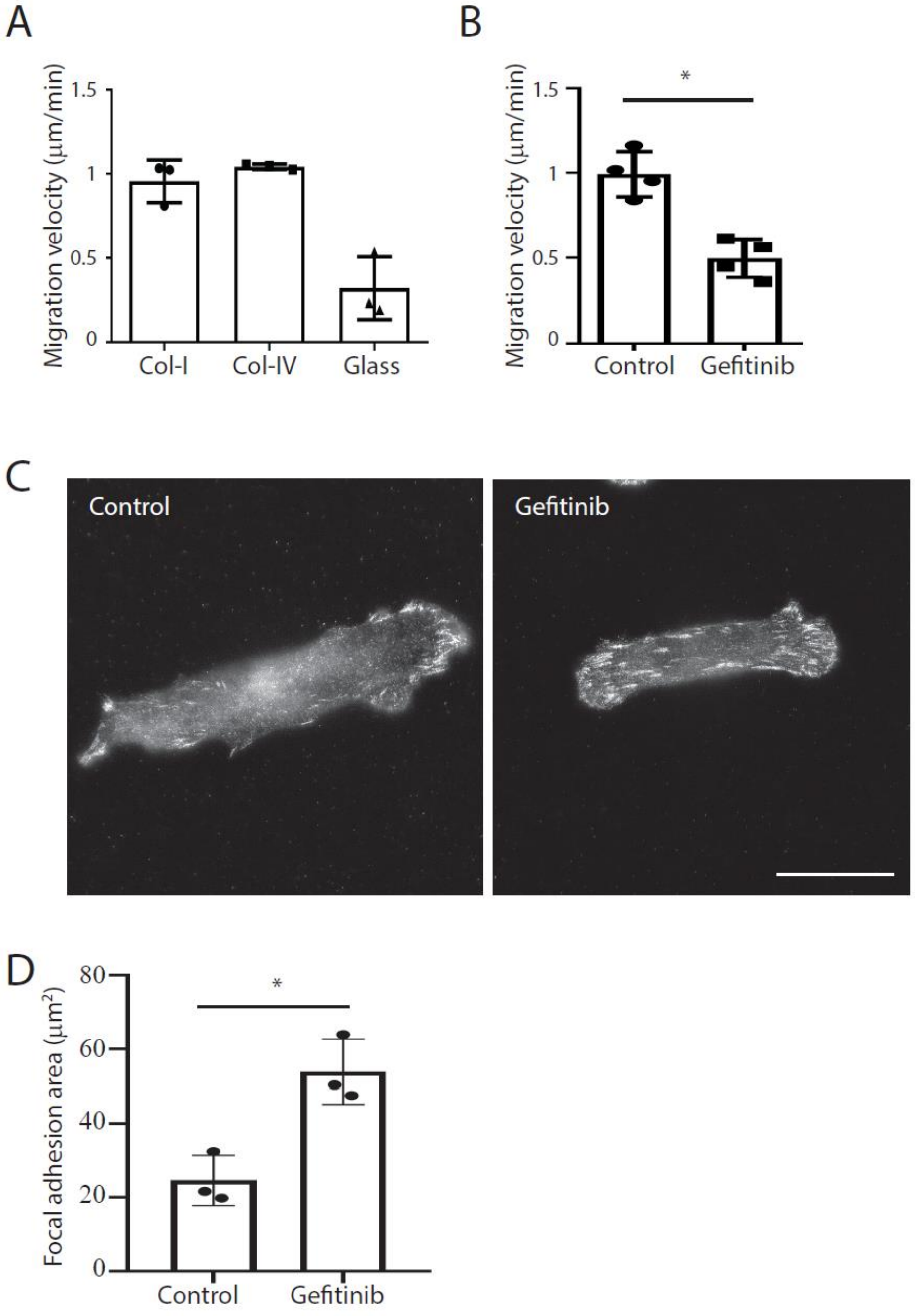
A
B
Migration velocity (μm/min)
1.5
1
0.5
0
Col-I
Col-IV
Glass
*
Control
Gefitinib
C
Control
Gefitinib
D
Focal adhesion area (μm²)
80
60
40
20
0
*
Control
Gefitinib

Supp. Fig. 1

Supp. Figure 1. **Characterization of cell response to collagen-I and to Gefitinib. a**, MDA-MB-231 cells were seeded on naked glass or collagen-I- or collagen-IV-coated glass and imaged every 10 min for 16h. Results are expressed as the mean migration velocity in µm/min. **b**, MDA-MB-231 cells treated or not with Gefitinib and seeded on collagen-I were imaged every 10 min for 16h. Results are expressed as the mean migration velocity in µm/min (*P<0.001, Student's t test. N=4). **c**, MDA-MB-231 cells seeded on collagen-I and treated or not with Gefitinib, as indicated, were fixed and stained for vinculin. Scale bar: 10 µm. **d**, Quantification of total focal adhesion area as in c (* P<0.001, Student's t test. N=3).

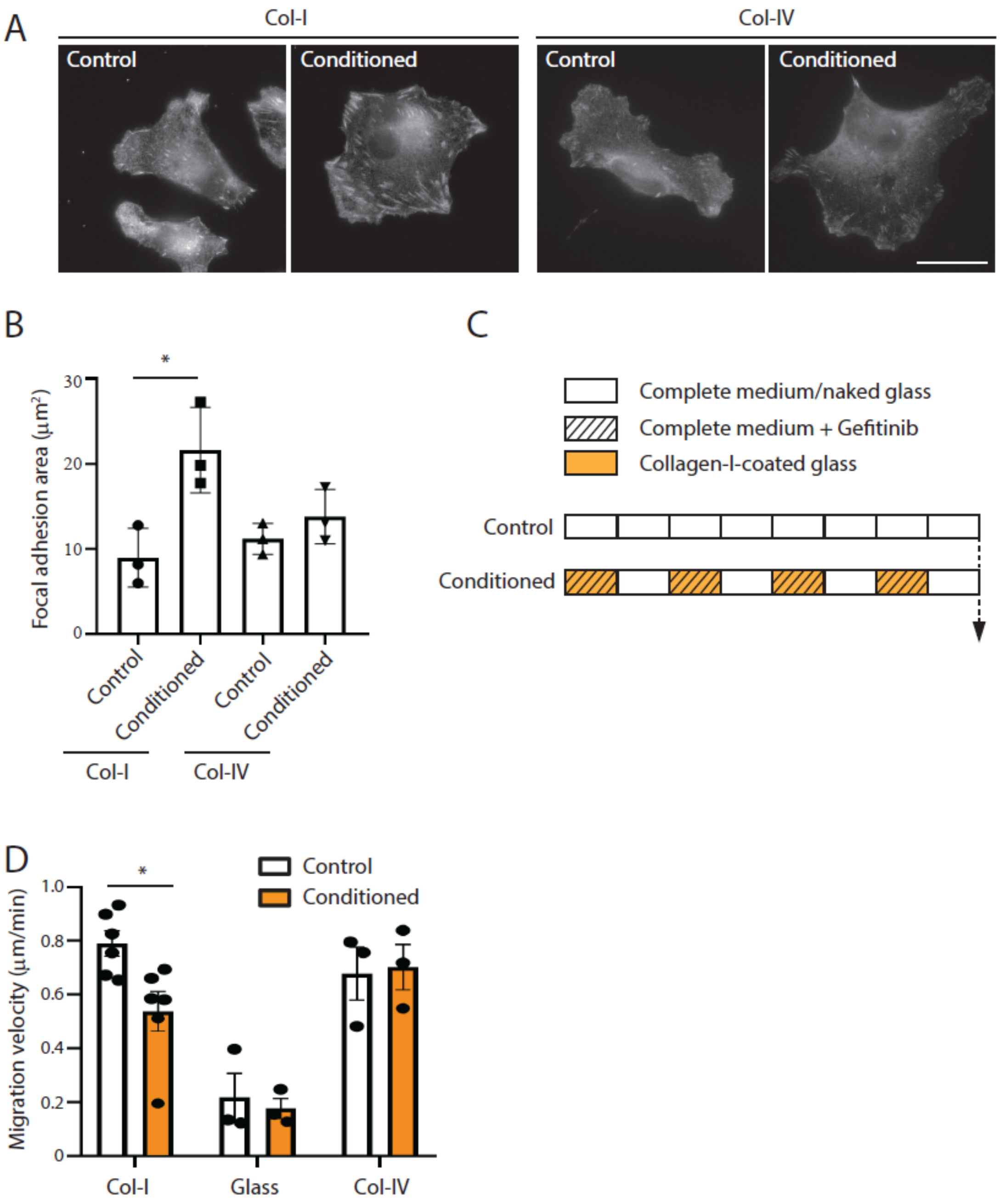
A
Col-I
Col-IV
Control
Conditioned
Control
Conditioned
B
Focal adhesion area (µm²)
30
20
10
0
*
Control
Conditioned
Control
Conditioned
Col-I
Col-IV
C
Complete medium/naked glass
Complete medium + Gefitinib
Collagen-I-coated glass
Control
Conditioned
D
Migration velocity (µm/min)
1.0
0.8
0.6
0.4
0.2
0
*
Control
Conditioned
Col-I
Glass
Col-IV

Supp. Fig. 2

Supp. Figure 2. **Characterization of the conditioned response in MDA-MB-231 and PC-9 cells. a**, Control or conditioned MDA-MB-231 cells were seeded on collagen-I or collagen-IV, as indicated, before to be fixed and stained for vinculin. Scale bar: 10 μm. **d**, Quantification of total focal adhesion area as in a (* P<0.001, One Way Analysis of Variance – ANOVA. N=3). **c**, Description of PC-9 cells culture conditions for the conditioning protocol using the collagen-I/Gefitinib association and for control cells. Cells were split every 3 days (vertical bars). **d**, Control or conditioned PC-9 cells seeded on the indicated extracellular matrix, were imaged every 10 min for 16h. Results are expressed as the mean migration velocity in μm/min (*P<0.001, One Way Analysis of Variance – ANOVA. N=3).

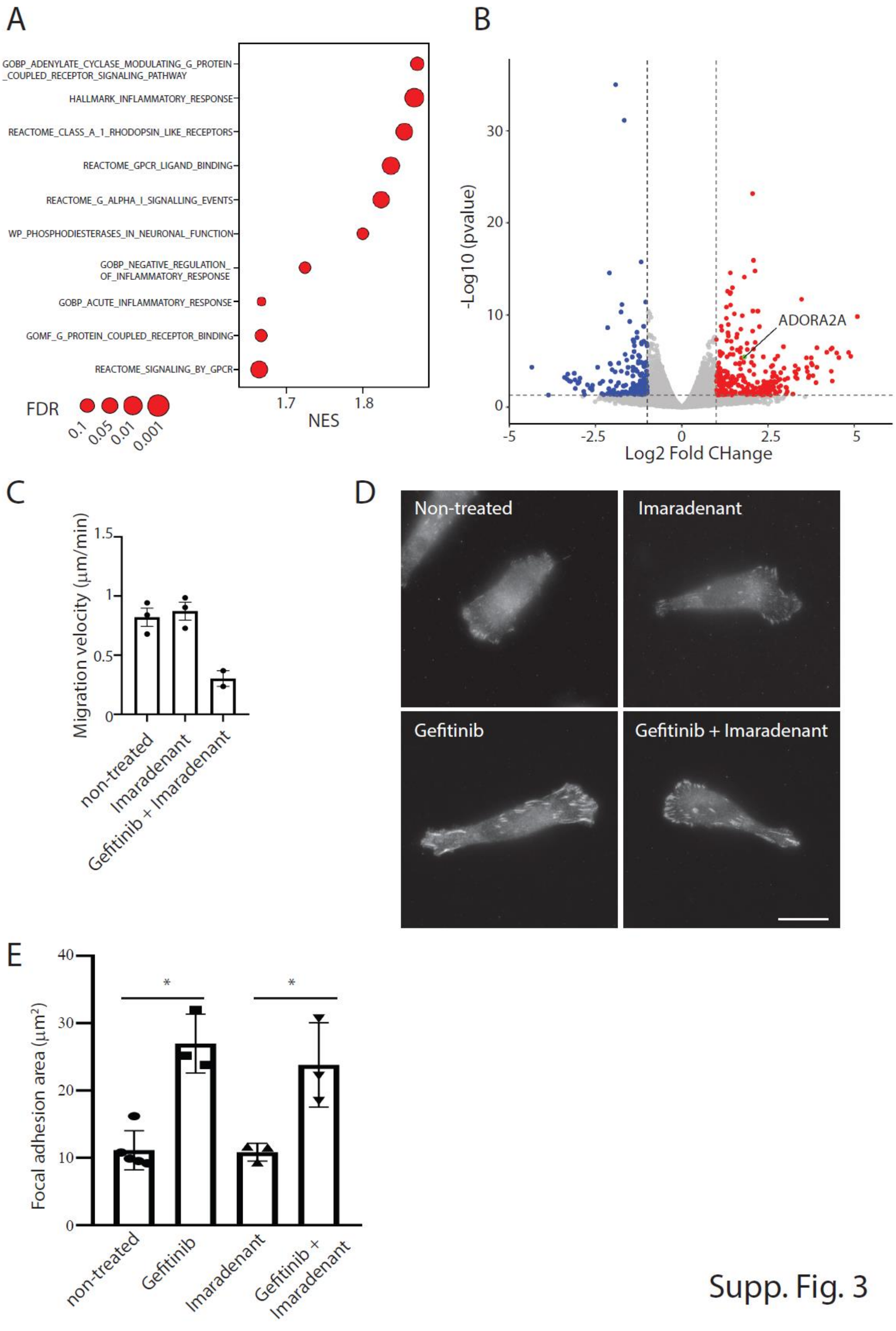

A
GOBP_ADENYLATE_CYCLASE_MODULATING_G_PROTEIN_COUPLED_RECEPTOR_SIGNALING_PATHWAY
HALLMARK_INFLAMMATORY_RESPONSE
REACTOME_CLASS_A_1_RHODOPSIN_LIKE_RECEPTORS
REACTOME_GPCR_LIGAND_BINDING
REACTOME_G_ALPHA_I_SIGNALLING_EVENTS
WP_PHOSPHODIESTERASES_IN_NEURONAL_FUNCTION
GOBP_NEGATIVE_REGULATION_OF_INFLAMMATORY_RESPONSE
GOBP_ACUTE_INFLAMMATORY_RESPONSE
GOMF_G_PROTEIN_COUPLED_RECEPTOR_BINDING
REACTOME_SIGNALING_BY_GPCR
FDR
0.1
0.05
0.01
0.001
1.7
1.8
NES
B
-Log10 (pvalue)
ADORA2A
Log2 Fold CHange
C
Migration velocity (µm/min)
non-treated
Imaradenant
Gefitinib + Imaradenant
D
Non-treated
Imaradenant
Gefitinib
Gefitinib + Imaradenant
E
Focal adhesion area (µm²)
non-treated
Gefitinib
Imaradenant
Gefitinib + Imaradenant


Supp. Fig. 3

Supp. Figure 3. **RNAseq and characterization of cell response to imaradenant. a**, List of enriched pathways using Gene Set Enrichment Analysis comparing control and conditioned MDA-MB-231 cells. **b**, Volcano plot of differentially expressed genes between control and conditioned MDA-MB-231 cells. **c**, MDA-MB-231 cells treated or not with imaradenant and Gefitinib, as indicated, and seeded on collagen-I were imaged every 10 min for 16h. Results are expressed as the mean migration velocity in μm/min. **d**, MDA-MB-231 cells seeded on collagen-I and treated or not with Gefitinib and imaradenant, as indicated, were fixed and stained for vinculin. Scale bar: 10 μm. **e**, Quantification of total focal adhesion area as in b (* $P<0.001$, One Way Analysis of Variance – ANOVA. N=3).

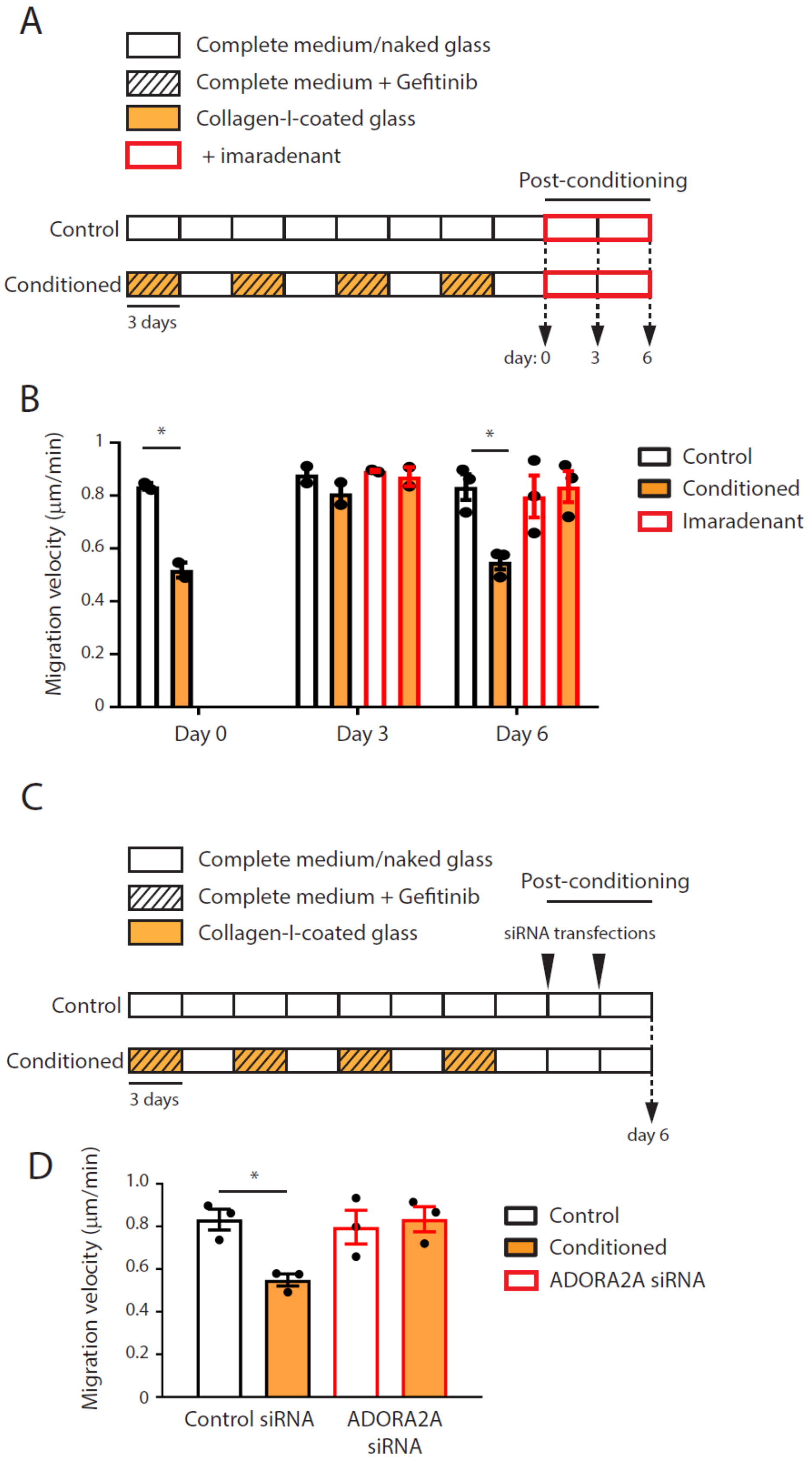
A
Complete medium/naked glass
Complete medium + Gefitinib
Collagen-I-coated glass
+ imaradenant
Post-conditioning
Control
Conditioned
3 days
day: 0
3
6
B
Migration velocity (µm/min)
1
0.8
0.6
0.4
0.2
0
*
*
Day 0
Day 3
Day 6
Control
Conditioned
Imaradenant
C
Complete medium/naked glass
Complete medium + Gefitinib
Collagen-I-coated glass
Post-conditioning
siRNA transfections
Control
Conditioned
3 days
day 6
D
Migration velocity (µm/min)
1.0
0.8
0.6
0.4
0.2
0
*
Control siRNA
ADORA2A siRNA
Control
Conditioned
ADORA2A siRNA

Supp. Fig. 4

Supp. Figure 4. **Role of ADORA2A in maintaining the conditioned response oscillation. a**, Description of MDA-MB-231 cells culture conditions for the conditioning and post-conditioning protocol using the collagen-I/Gefitinib association and for control cells. Cells were split every 3 days (vertical bars). **b**, Control or conditioned MDA-MB-231 cells kept on naked glass for the indicated time post-conditioning and treated or not with imaradenant, as indicated, were seeded on collagen-I and imaged every 10 min for 16h. Results are expressed as the mean migration velocity in μm/min (* P<0.001, One Way Analysis of Variance – ANOVA. N=2). **c**, Description of MDA-MB-231 cells culture conditions for the conditioning protocol and post-conditioning treatment with ADORA2A specific siRNAs. Cells were split every 3 days (vertical bars). **d**, Control or conditioned MDA-MB-231 cells treated as in c were seeded on collagen-I and imaged every 10 min for 16h. Results are expressed as the mean migration velocity in μm/min (* P<0.001, One Way Analysis of Variance – ANOVA. N=3).

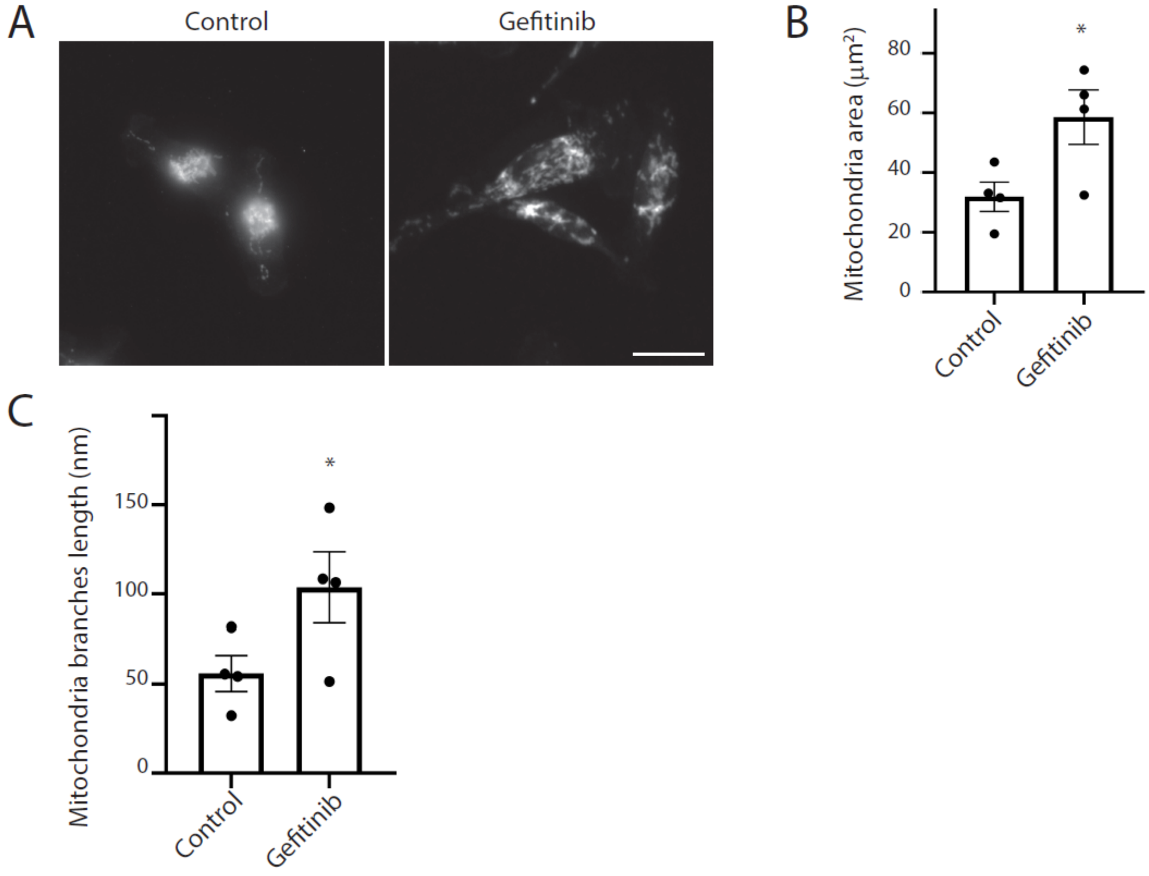
A
Control
Gefitinib
B
Mitochondria area (μm²)
80
60
40
20
0
*
Control
Gefitinib
C
Mitochondria branches length (nm)
150
100
50
0
*
Control
Gefitinib

Supp. Fig. 5

Supp. Figure 5. **Characterization of mitochondria dynamics. a**, Control MDA-MB-231 cells seeded on collagen-I and treated or not for 24h with Gefitinib were fixed and stained for Tomm20. Scale bar: 10 μm. **b-c**, Quantification of total mitochondria area (b) and of the total length of branches (c) in cells as in a (* $P<0.001$, Student's t test. $N=4$).
